# Individual response to mTOR inhibition in delaying replicative senescence of mesenchymal stromal cells

**DOI:** 10.1101/420026

**Authors:** Eliane Antonioli, Natália Torres, Mario Ferretti, Carla A. Piccinato, Andrea L. Sertie

**Author notes:** Corresponding author: (ALS).

## Abstract

**Background aims:** Delaying replicative senescence and extending lifespan of human mesenchymal stromal cells (MSCs) may enhance their potential for tissue engineering and cell based therapies. Accumulating evidence suggests that inhibitors of the mTOR signaling pathway, such as rapamycin, constitute promising pharmacological agents to retard senescence and extend stemness properties of various progenitor cell types. Here, we investigated whether the ability of rapamycin to postpone replicative senescence varies among bone marrow MSC samples (BM-MSCs) derived from different healthy donors, and explored the molecular mechanisms that drive rapamycin-mediated lifespan increment.

**Methods:** BM-MSCs at early passages were serially passaged either in absence or continuous presence of rapamycin and the number of cell population doublings until growth arrest was measured. The inhibition of mTOR signaling was assessed by the phosphorylation status of the downstream target RPS6. The expression levels of several senescence and pluripotency markers at early and late/senescent passages were analyzed by RT-qPCR, flow cytometry and western blot.

**Results:** We found that the lifespan extension in response to the continuous rapamycin treatment was highly variable among samples, but effective in most BM-MSCs. Despite all rapamycin-treated cells secreted significantly reduced levels of IL6, a major SASP cytokine, and expressed significantly higher levels of the pluripotency marker *NANOG*, the expression patterns of these markers were not correlated with the rapamycin-mediated increase in lifespan. Interestingly, rapamycin-mediated life-span extension was significantly associated only with repression of p16^INK4A^ protein accumulation.

**Conclusions:** Taken together, our results suggest that some, but not all, BM-MSC samples would benefit from using rapamycin to postpone replicative arrest and reinforce a critical role of p16^INK4A^ protein downregulation in this process.

## Introduction

In recent years, there has been increasing evidence that persistent activation of the growth-promoting mammalian target of rapamycin (mTOR) pathway plays a central role in cellular senescence and organismal aging [1–3], representing a key molecular driver of stem cell depletion and reduced tissue regenerative capacity [4–6]. Importantly, attenuation of mTOR signaling with rapamycin seems to preserve the clonogenic ability and function, besides delaying the activation of senescence mechanisms, in mouse and human stem cells from various tissues, including hematopoietic [5, 7, 8], epithelial [6, 9], spermatogonial [10] and mesenchymal stem cells [11].

The mechanisms by which mTOR inhibition protects from stem cell exhaustion and aging have been associated with reduced accumulation of cytoplasmic and/or mitochondrial reactive oxygen species [6, 7, 11] and DNA damage [6, 9, 11], decreased secretion of major senescence-associated cytokines [6] and reduced expression of tumor suppressors such as p16^INK4A^ [6, 11]. Moreover, the rapamycin-induced persistent expression of pluripotency genes, such as *NANOG* and *OCT4* [11], has been suggested to contribute to the retention of stemness properties of stem cells.

The possibility of retarding senescence and extending stemness properties of *in vitro* expanded mesenchymal stromal cells (MSCs) is particularly relevant to regenerative medicine, as replicative senescence might limit the number of cells required for clinical use [12, 13]. In a previous study, using long-term MSC cultures, we have shown that bone marrow MSCs (BM-MSCs) isolated from healthy young donors display variable *in vitro* replicative potential until reaching senescence and ceasing to proliferate [14]. Also, we documented that those BM-MSC samples with lower expression of the senescence marker p16^INK4A^ and higher expression of the pluripotency marker *OCT4* at early passages present greater replicative lifespan [14]. Although rapamycin has been shown to decelerate cell senescence in different experimental models, such as radiation and replicative induced senescence, no study has evaluated the effects of long-term treatment of BM-MSC samples endowed with variable replication capabilities with rapamycin. These observations prompted us to ask whether the ability of rapamycin to postpone replicative senescence varies among individual BM-MSC samples and to investigate the molecular players involved in lifespan extension mediated by mTOR inhibition in this long-term cell culture model.

## Materials and methods

### Cell culture and long-term inhibition of mTOR (rapamycin treatment)

Primary human BM-MSCs of five healthy young adults (3 males and 2 females, aging 30-39 years old) have been previously isolated and characterized [14]. The samples - referred to as BM09, BM12, BM13, BM16 and BM18 - were taken after written consent from donors, and the study was approved by the Ethics Committee of Hospital Israelita Albert Einstein.

Cells at an early passage (passage 5) were thawed and cultured under standard conditions as monolayers in DMEM-low glucose (Thermo Fisher Scientific, cat. 31600-034) supplemented with 15% fetal bovine serum (FBS, Thermo Fisher Scientific, cat. 12483-020), 1 mM L-glutamine (Thermo Fisher Scientific, cat. 25030081) and 1% antibiotic-antimycotic solution (Thermo Fisher Scientific, cat. 15240-062) in T-25 flasks at 37°C in a humidified atmosphere containing 5% CO_2_. In order to inhibit mTOR signaling, rapamycin (Sigma Aldrich, cat. R0395) was used at a final concentration of 20nM based both on previous studies [6, 9] and on pilot dose-response studies of our group that have shown that either 20nM or 50nM of rapamycin were able to almost completely inhibit mTOR signaling, while maintaining the proliferative capacity of the cells (data not shown). Cells, cultured with either rapamycin or DMSO (Sigma, cat. D2650; used as vehicle control), were serially passaged at a density of 4000 cells/cm^2^ every 7 days until ceasing to proliferate (becoming senescent). Culture media (with and without rapamycin) were changed every two days. The number of cell population doublings in both conditions was assessed by the Trypan Blue exclusion method.

Cumulative cell population doublings (PD) in each conditions (with and without rapamycin) was calculated using the following equation: log_10_(NH/N1)/log^10^(2), where NH= cell harvest number and NI= plating cell number. The population doubling time (PDT) was calculated as follows: log_10_(2)×T_H-I_/[log_10_(*N*_H_/*N*_I_)], where T_H–I_ is time between harvest and inoculum.

To examine whether the observed lifespan extension is rapamycin-dependent, rapamycin was withdrawn from BM09 cells during the exponential phase of growth and cells were cultured in rapamycin-free medium until proliferative arrest. After this, quiescent cells were cultured again in culture medium containing rapamycin (20nM). Results shown are representative of at least two independent experiments.

### Immunoblotting

Total protein extracts from rapamycin-treated and untreated (DMSO) cells were obtained at passages when untreated cells entered senescence and stop proliferating and at passages when rapamycin-treated cells entered into proliferative arrest using Ripa Buffer (Sigma, cat. R0278) containing protease and phosphatase inhibitor cocktails (Sigma Aldrich, cat. P8340 and P5726). Equal amounts (20μg) of proteins were separated by SDS-PAGE and transferred to nitrocellulose membranes, which were then blocked and incubated with the following primary antibodies: anti-P16^INK4A^ (Abcam, ab108349), anti-phospho-RPS6^S240/244^ (Cell Signaling Technology, #5364), and ß-actin (Sigma, A2228) for loading control. Detection was performed using horseradish peroxidase-coupled anti-rabbit or anti-mouse secondary antibodies (Cell Signaling Technology, #7074 and #7076), ECL substrate (GE Healthcare, cat. RPN2236), and the ChemiDoc™ MP Imaging System (Bio-Rad). The intensity of the bands was determined by densitometry using The Image Lab Software (Bio-Rad). The immunoblotting experiments were repeated more than three times and similar results were observed.

### Measurement of inflammatory cytokines

The levels of proinflammatory cytokines IL6 and IL8 in culture supernatants from rapamycin-treated and untreated (DMSO) cells at passages when untreated cells have entered proliferative arrest were evaluated by flow cytometry using the Cytometric Bead Array (CBA) Human Inflammatory Cytokine Kit (BD Biosciences, cat. 551811) following the manufacturer’s instructions. For data analysis, FCAP Array software (Soft Flow Inc.) was used. For each sample, secreted cytokine levels were normalized to cell number and expressed as pg/mL/cells. Results shown are representative of at least two independent experiments.

### Gene Expression Analyses (RT-qPCR)

Total RNA was extracted from rapamycin-treated and untreated (DMSO) cells at passages when untreated cells entered proliferative senescence using the Illustra RNAspin Mini RNA Isolation Kit (GE Healthcare, cat. 25-0500-71). RNA quality and quantity was assessed using the NanoDrop™ 3300 Fluorospectrometer (Thermo Fisher Scientific). The reverse transcription of 1 µg of total RNA was performed with QuantiTect Reverse Transcription kit (QIAGEN, cat. 205311). The qPCR were carried out using gene-specific primers (*OCT4, NANOG*, and *SOX2*) and the Maxima SYBR Green qPCR Master Mix (Thermo Fisher Scientific, cat. 0221) or the p16^INK4A^/CDKN2A predesigned TaqMan gene expression assay (Hs99999189_m1, Thermo Fisher Scientific) in an ABI 7500 real-time PCR system (Applied Biosystems), according to the manufacturer’s instruction. The expression levels of the target genes were normalized to the *GAPDH* housekeeping gene. Primer sequences used for qPCR were described previously [14]. All reactions were performed in triplicate. Results are expressed as the mean fold change of the normalized gene expression relative to a calibrator sample (#636690 reference RNA for RT-qPCR, Clontech) using the comparative CT method (^ΔΔ^Ct method). The RT-qPCR results are representative of two independent experiments.

### Statistical analysis

Statistical analyses were carried out using the SAS statistical analysis program (Statistical Analysis System Institute Inc., Cary, NC, USA). All correlation analyses were performed by the CORR procedure from at least duplicated results using the Spearman correlation method. The means obtained were calculated by the PROC GLM procedures of SAS and for that, log transformation was applied as needed. In all analysis, the level of significance was considered when p<0.05.

## Results

### MSCs from different donors exhibit variable lifespan extension in response to continuous mTOR inhibition

To evaluate the effects of mTOR inhibition on lifespan extension of BM-MSC samples derived from 5 healthy young donors (referred to as BM09, BM12, BM13, BM16 and BM18), which were previously shown to display high heterogeneity in their proliferative capacity [14], we cultivated these cells and serially passaged them in the same growth medium supplemented or not with rapamycin during the entire replicative lifespan, and the number of cumulative cell population doublings (PDs) and PD time (PDT) until cell cycle arrest were measured in both conditions (rapamycin-treated and untreated conditions).

First, we observed that rapamycin delayed the development of senescence-associated phenotype as all cell samples expanded in the presence of rapamycin displayed a more elongated spindle-like shape during almost the entire replicative lifespan, whereas the corresponding untreated cells assumed the enlarged senescence-associated morphology at relatively early passages. Next, we observed that BM-MSCs from different donors presented variable lifespan extension in response to the continuous presence of rapamycin: while rapamycin delayed replicative senescence and extended dramatically the lifespan of 1 sample (BM09: 23 additional PDs compared with the corresponding untreated cells), it had a moderate impact on serial expansion of 3 samples (BM18: 7 additional PDs, BM13: 5 additional PDs and BM16: 3 additional PDs compared with the corresponding untreated cells), and no impact on another sample (BM12) (Fig 1A). Also, we observed that the PDT of most samples treated with rapamycin remains significantly more constant for a longer period over the course of serial passages compared to the corresponding untreated cells, and this effect is most evident for BM09, which showed the greater lifespan extension promoted by rapamycin (Fig 1B). This important effect on BM09 lifespan was mediated by the continuous presence of rapamycin, as these cells lose their proliferative potential few passages after rapamycin removal and returned to active proliferation after replacement of the drug in culture medium (Fig 1C), suggesting that the cells achieved the quiescent state and retained the capacity to resume proliferation upon mTOR inhibition.

**Fig 1.**
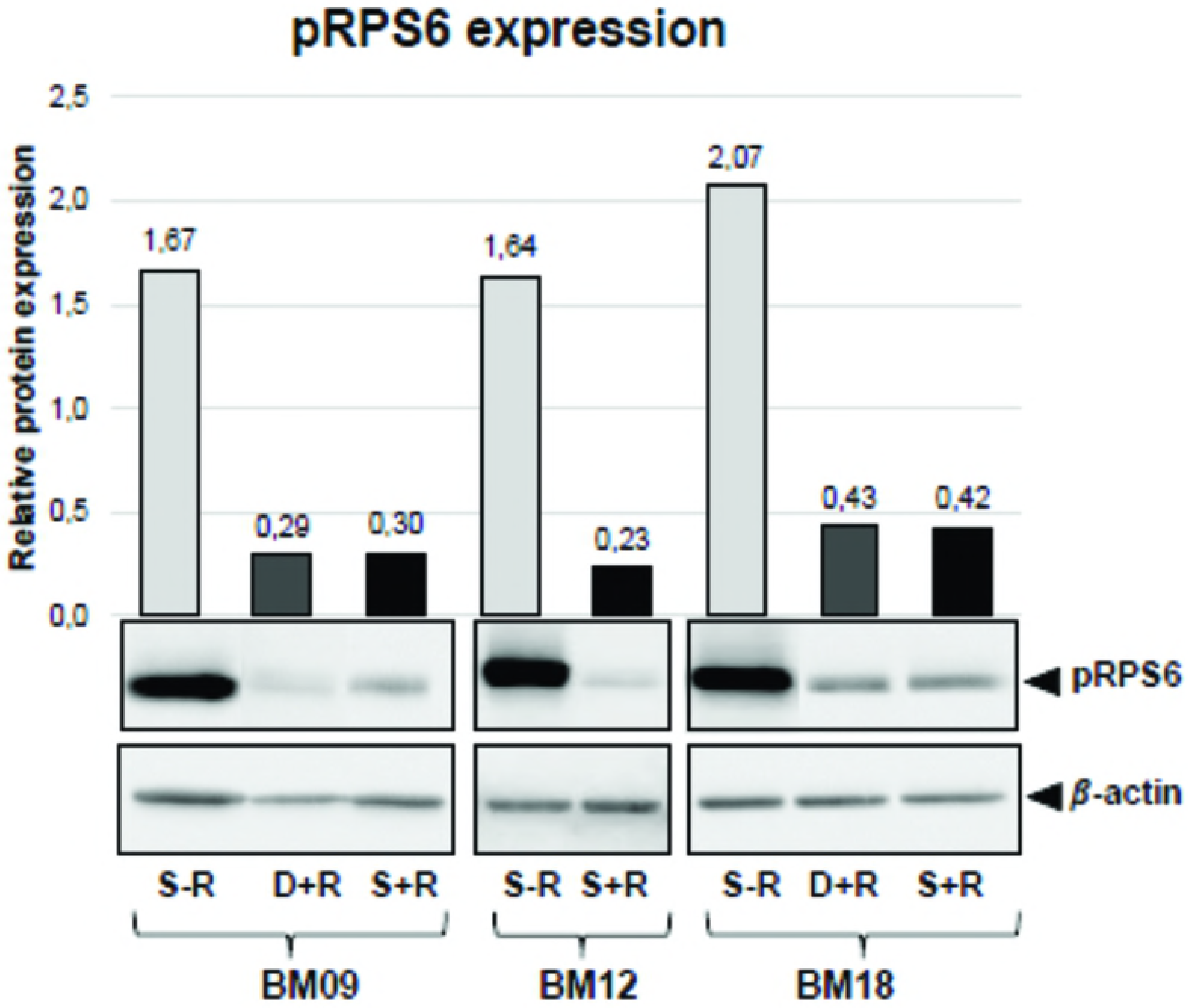
Lifespan extension and growth kinetics in response to continuous mTOR inhibition varies among different BM-MSC samples. (A) Cumulative population doubling (PD) curves of BM-MSC samples derived from 5 healthy young donors (BM09, BM12, BM13, BM16 and BM18) until replicative arrest in control conditions (DMSO) or in the continuous presence of rapamycin (RAPA). Each symbol represents a passage of rapamycin-treated (triangle) and untreated (dot) cells. The passage when untreated BM09, BM13, BM16 and BM18 samples entered replicative senescence while the corresponding rapamycin-treated cells continue to proliferate is referred to as the “deviation passage” and is indicated by an arrow. Since rapamycin had no impact on lifespan extension of BM12, no “deviation passage” was assigned for this sample. (B) PD time (PDT) of BM-MSC samples at each passage in control conditions (dot) or in the continuous presence of rapamycin (triangle). (C) Cumulative PD curves of BM09 in which rapamycin was removed (RAPA removal) until cells ceased growth and then replaced in culture medium (RAPA replacement). Each dot represents a passage of cells. Data shown in panels A to C are representative of results from at least two independent experiments.

Finally, we observed that despite the fact that all rapamycin-treated samples progress towards growth arrest, cells continued to respond to rapamycin along the entire lifespan, as assessed by the phosphorylation status of ribosomal protein S6 (RPS6), a downstream target of the mTOR pathway (Fig 2), which may suggest that mTOR-independent mechanisms also may contribute to replicative senescence. As depicted in Fig 2 and described hereafter, the passage when untreated cells entered replicative senescence is referred to as the “senescent passage without rapamycin”, the same passage of the corresponding rapamycin-treated samples (passage in which most samples still displayed proliferation ability) is referred to as the “deviation passage with rapamycin”, and the passage when rapamycin-treated cells entered into proliferative arrest is referred to as “senescent passage with rapamycin”.

**Fig 2.**
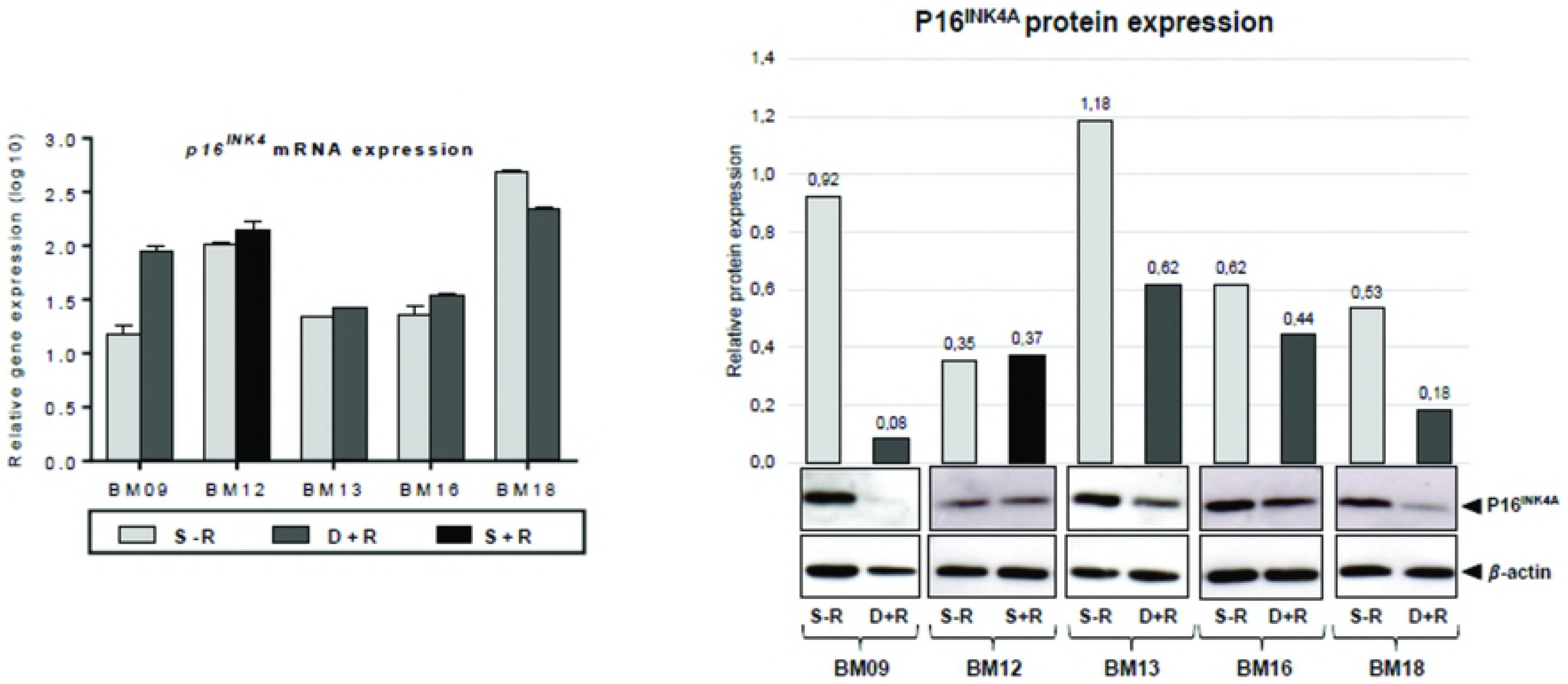
The ability of BM-MSCs to respond to rapamycin continues along the entire replicative lifespan. mTOR signaling inhibition is reflected by the phosphorylation status of RPS6, a downstream target of the mTOR pathway. Phosphorylated RPS6 (pRPS6) was quantified by western blot in untreated and rapamycin-treated BM09 and BM18 samples at passages when untreated cells entered replicative arrest (S-R= senescent passage without rapamycin; D+R= deviation passage with rapamycin), as well as when rapamycin-treated cells stop proliferating (S+R= senescent passage with rapamycin). Since rapamycin had no impact on lifespan extension of BM12, pRPS6 levels were measured in untreated and rapamycin-treated cells at the same senescent passage. Band intensities were densitometrically evaluated and bar graphs above bands represent the densitometric values of pRPS6 normalized to the loading control (*β*-actin). pRPS6 levels were greatly reduced by continuous rapamycin treatment. No consistent differences were seen in pRPS6 levels between rapamycin-treated BM09 and BM18 cells at the deviation passage (D+R) and at the final senescent passage (S+R). Data show representative blots from at least two independent experiments.

### Expression levels of known senescence and pluripotency markers in early-passage MSCs have no association with lifespan extension promoted by continuous mTOR signaling attenuation

We next evaluated whether the expression levels of senescence-associated proteins (cytoplasmic proteins p16^INK4A^, p21^WAF1^, pRPS6 and SOD2, as well as secreted SASP cytokines IL6 and IL8) and pluripotency-related genes (*NANOG, SOX2* and *OCT4*, whose protein expression in MSCs is rather low and poorly detected by Western blot) in all BM-MSC samples at early passages (5^th^ or 6^th^ passages) which we have previously quantified [14], could predict an individual lifespan extension in response to the continuous presence of rapamycin. Here we observed that the expression levels of these markers at early passages were not significantly correlated with the rapamycin-mediated increase in the replicative capacity of each sample expanded in normal medium (p16^INK4A^ r= 0.20 p= 0.78; p21 ^WAF1^ r=-0.60 p=0.35; pRPS6 r=-0.50 p=0.45; SOD2 r=0.30 p=0.68; IL6 r=-0.20 p=0.78; IL8 r=0.30 p=0.68; *NANOG* r=0.20 p=0.78; *SOX2* r=-0.40 p=0.51; *OCT4* r=-0.10 p=0.95) (Supplementary Figure 1a). Additionally, we did not observe any significant correlation between the inherent replicative capacity of untreated BM-MSC samples and the extent of lifespan extension promoted by rapamycin (r=0.20, p=0.78); in other words, the replicative potential (number of PD until cell cycle arrest) of untreated BM-MSC samples do not predict their response to rapamycin (S1 Fig).

### mTOR attenuation significantly reduced the secretion of the SASP protein IL6 and increased the expression of the pluripotency gene NANOG in MSCs

To verify whether rapamycin treatment in our long-term BM-MSC culture model leads to the regulation of molecular events often associated with stem cell senescence, we analyzed the gene and protein expression of p16^INK4A^, the secretion of cytokines IL6 and IL8, and the expression of pluripotency genes *OCT4* and *NANOG* in rapamycin-treated and untreated cells at passages when untreated cells entered into a state of proliferative arrest (referred to as the “deviation passage”, as depicted in Fig 1).

We observed that whereas rapamycin exerted no significant effect on p16^INK4A^ mRNA expression, it decreased p16^INK4A^ protein expression in BM09, BM13, BM16 and BM18, but had no effect on the expression of this protein in BM12 (Fig 3A). Therefore, due to the lack of effect of mTOR inhibition on p16^INK4A^ protein expression in BM12, there was not a significant effect of rapamycin treatment on p16^INK4A^ levels when cells are analyzed in group (rapamycin-treated versus untreated cells; p=0.06).

**Fig 3.**
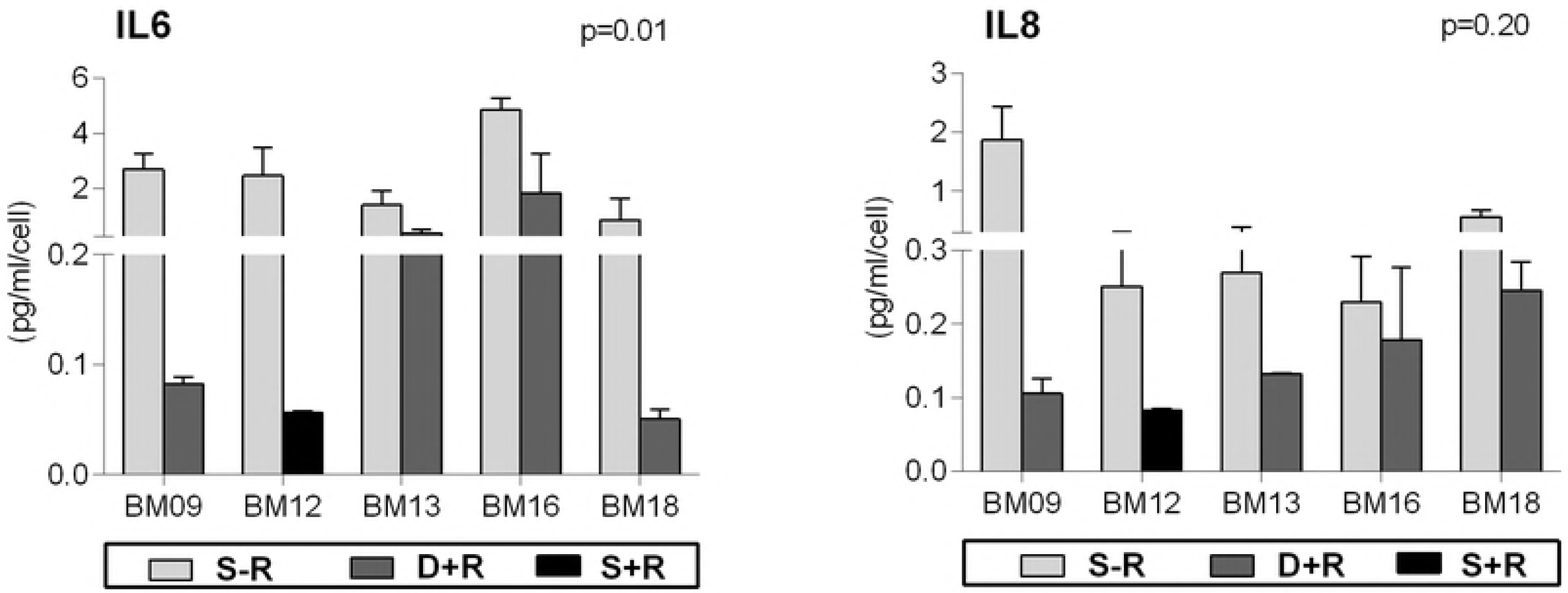
mTOR signaling inhibition leads to the regulation of molecular events often associated with stem cell senescence. The expression of senescence- and pluripotency-related markers were analyzed in untreated and rapamycin-treated BM09, BM13, BM16 and BM18 samples at passages when untreated cells entered replicative senescence (deviation passage). Since rapamycin had no impact on lifespan extension of BM12, the expression levels of these markers were measured in untreated and rapamycin-treated cells at the same senescent passage. (A) p16^INK4A^ gene expression levels were analyzed by RT-qPCR. The bar graphs show relative gene expression levels after normalization to *GAPDH*. p16^INK4A^ protein expression levels were analyzed by western blot. Band intensities were densitometrically evaluated and bar graphs above bands represent the densitometric values of p16^INK4A^ normalized to the loading control (β-actin). These results are representative of at least two independent experiments. (B) The levels of IL6 and IL8 secretion were determined using a CBA proinflammatory kit. The bar graphs represent the obtained concentration values for IL6 and IL8 (pg/mL) normalized against cell numbers. These results are representative of two independent experiments performed in triplicate. (C) The gene expression levels of *OCT4, SOX2* and *NANOG* were analyzed by RT-qPCR. The bar graphs show relative gene expression levels after normalization to *GAPDH*. These results are representative of two independent experiments performed in triplicate. S-R= senescent passage without rapamycin; D+R= deviation passage with rapamycin; S+R= senescent passage with rapamycin.

Although all rapamycin-treated samples showed, as expected, decreased secretion of IL6 and IL8 compared to the corresponding untreated cells, only the effect of mTOR inhibition on IL6 secretion reached statistical significance (p<0.05) (Fig 3B). This lack of significance may be probably due to the highly variable IL8 secretion among samples. Finally, we observed that cells expanded in the presence of rapamycin showed significant increased expression levels of *NANOG* (p<0.05) (Fig 3C). The rapamycin-mediated significant decrease in the secretion of a major SASP cytokine and increase in the expression of *NANOG* may contribute to cell proliferation at a relatively more constant rate (less variable PDT, Fig 1B) over an extended period in culture and retention of the non-senescent phenotype of MSCs.

### Extension of MSCs lifespan by continuous attenuation of mTOR signaling is correlated with downregulation of p16^INK4A^ protein expression

We then evaluated whether the rapamycin-mediated decrease in the expression of senescence-associated markers p16^INK4A^, IL6 and IL8 and increase in the expression of pluripotency gene *NANOG* at the “deviation passage” are correlated with the rapamycin-mediated lifespan extension. We observed that only p16^INK4A^ expression was significantly and inversely correlated with lifespan promoted by mTOR signaling inhibition (r=-1.0, p=0.016), as the greater the fold reduction in p16^INK4A^ expression levels, the greater the rapamycin extended lifespan (Fig 4). These results suggest that the ability of rapamycin in inhibiting p16^INK4A^ accumulation, rather than inhibiting IL6 and IL8 secretion and stimulating *NANOG* expression, during serial passages is tightly associated with the MSC replicative lifespan extension.

**Fig 4.**
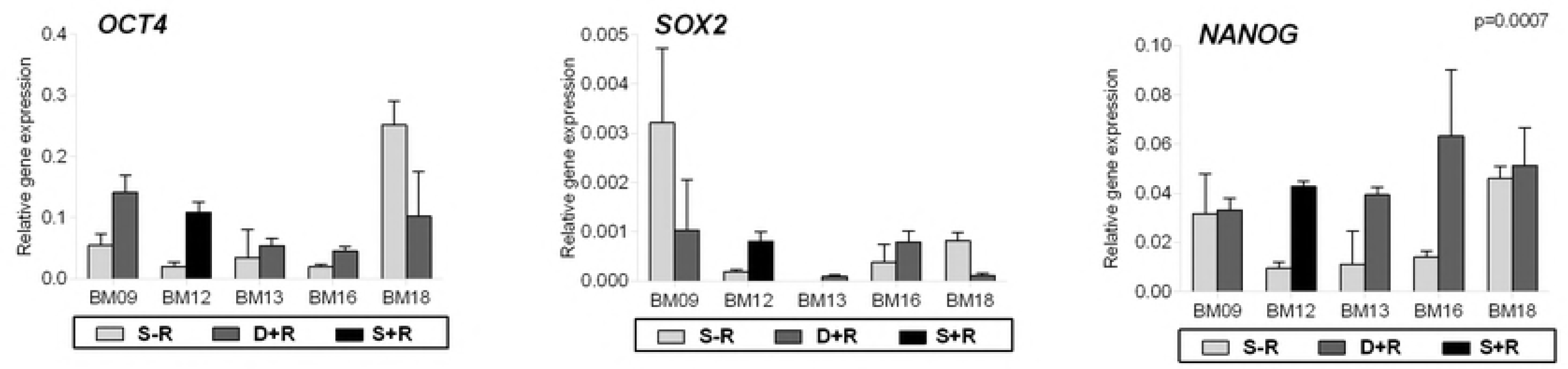
Downregulation of p16^INK4A^ expression is correlated with lifespan extension promoted by the continuous presence of rapamycin. The fold reduction in p16^INK4A^ protein expression and in IL6 and IL8 secretion, and the fold increase in *NANOG* gene expression between untreated and rapamycin-treated BM09, BM13, BM16 and BM18 samples at passages when untreated cells entered replicative senescence (deviation passage), or in the case of BM12 at the same senescence passage, were plotted against the additional PD number obtained for the corresponding rapamycin-treated cells and statistically analyzed by Spearman correlation, as shown in the graphs.

Finally, since rapamycin-treated BM-MSCs reached proliferative arrest, we sought to verify whether the amount of p16^INK4A^ protein is maintained at lower levels until the final passage or it is accumulated again in the arrested cells. We observed that the expression levels of p16^INK4A^ in rapamycin-treated cells that ceased to proliferate (at the “senescent passage with rapamycin”) are slightly greater than those observed in the “deviation passage with rapamycin” (Fig 5), suggesting that the slight increase in p16^INK4A^ accumulation may contribute to the cessation of rapamycin-treated cell proliferation, but that other molecular events might also be required to induce final growth arrest of rapamycin-treated cells.

**Fig 5.**
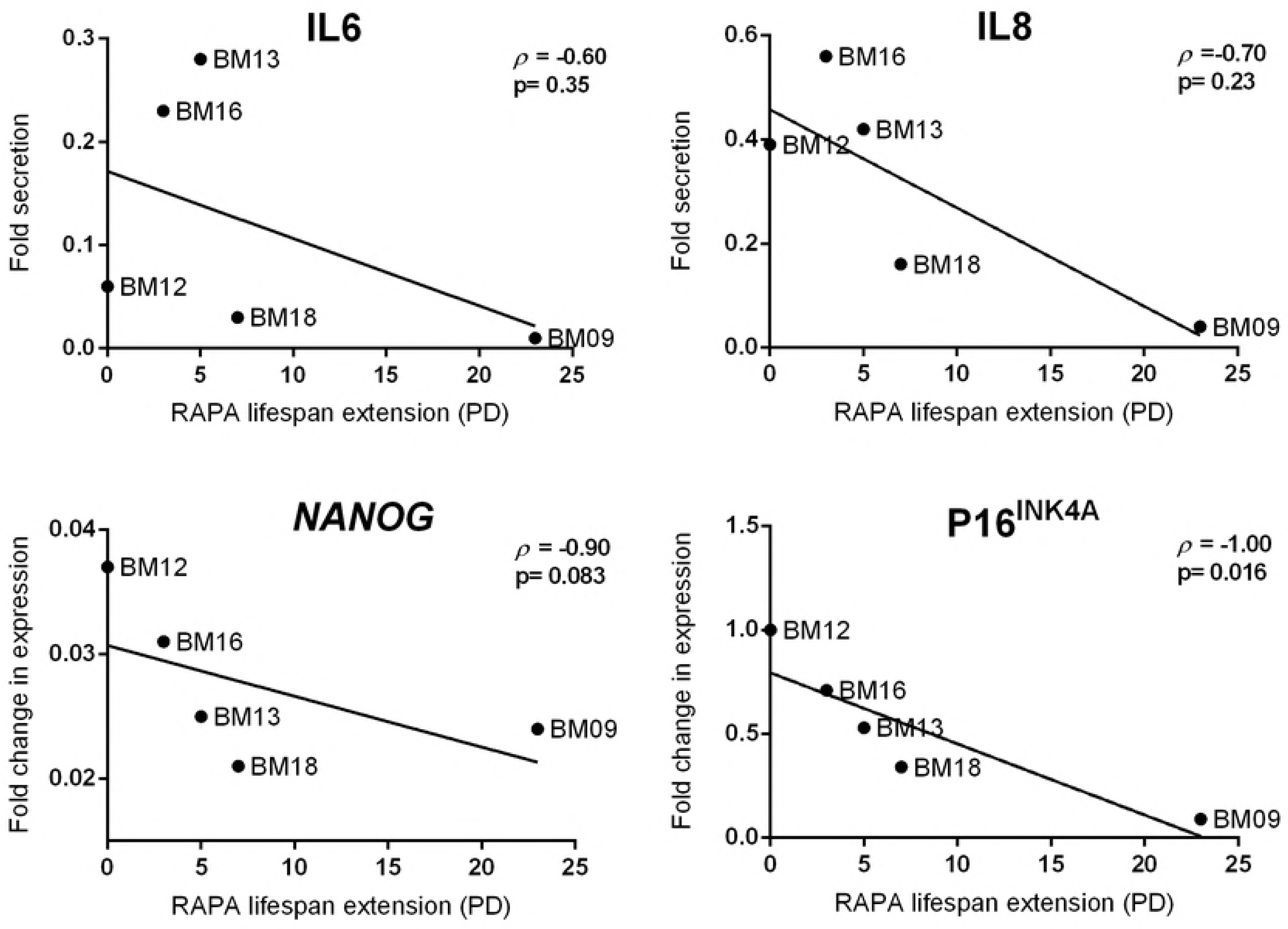
p16^INK4A^ protein expression increased slightly in rapamycin-treated cells that stopped proliferating. The protein expression levels of p16^INK4A^ were analyzed by western blot in rapamycin-treated BM-MSCs at the deviation passage with rapamycin (D+R) and at the senescent passage with rapamycin (S+R). Band intensities were densitometrically evaluated and bar graphs above bands represent the densitometric values of p16^INK4A^ normalized to the loading control (*β*-actin). The results obtained for BM12 is already showed in Fig 3a. Data show representative blots from at least two independent experiments.

## Discussion

It is well established that MSC cultures from a wide variety of healthy tissues have finite replicative lifespan and cease proliferating after a certain number of divisions [12]. The need for extensive MSC expansion for cellular therapy applications led to the search for new approaches to obtain sufficient number of cells before replicative senescence is reached. mTOR specific inhibitors, such as rapamycin, have emerged as potential adjuvant candidates for retarding the process of replicative senescence [15, 16]. Despite not yet being used for cellular therapy purposes, rapamycin has already been approved by the U.S. Food and Drug Administration (FDA) for a variety of clinical applications, including immunosuppressive and anticancer treatments [17].

Data from the present study show that rapamycin has variable ability of postponing replicative senescence of BM-MSC samples derived from different healthy donors. Interestingly, even cell samples that show greater inherent proliferation capabilities (BM09, BM12 and BM18) respond distinctly to rapamycin, despite that fact that no clinically relevant differences were observed among donors, and cells were expanded under the same standardized controlled conditions [14]. The effects of rapamycin in BM-MSC lifespan and aging was tested using a long-term *in vitro* BM-MSC expansion model, in which cells were continuously exposed to rapamycin treatment and exhibited attenuated mTOR signaling until proliferative arrest, as monitored by the phosphorylation status of RPS6, a key downstream protein target of mTOR pathway. Others studies, using distinct *in vitro* models of cellular senescence, have shown that mTOR inhibition by rapamycin can delay the progression of stem cell senescence [6, 11]. However, results of our study provide evidence for the benefits of mTOR inhibition in long-term MSC expansion and also, for the first time, alert for marked interindividual variability in the response to rapamycin, suggesting that its use might benefit expansion of some cells, not all BM-MSCs.

The identification of reliable early predictors of stem cell lifespan extension in response to mTOR inhibition would be valuable for regenerative medicine and cellular therapy. However, none of the six classical senescence-associated proteins and the three pluripotent-related genes expressed by early-passage BM-MSCs evaluated here were useful predictors of rapamycin-mediated lifespan prolongation, as no correlation could be established.

To characterize important players involved in rapamycin-mediated postponement of replicative senescence of BM-MSCs in our model, we analyzed the expression levels of known senescence- and pluripotency-related markers in rapamycin-treated and untreated cells at the passage when untreated cells stop proliferating (become senescent) and the corresponding rapamycin-treated cells still displayed proliferation capacity. Our results showed that, as observed in other models of cellular senescence, rapamycin-treated cells secreted significantly reduced levels of a major SASP cytokine (IL6), which is known to act in an autocrine manner to reinforce the cell cycle arrest of senescent cells [6, 18], and expressed significantly increased levels of *NANOG*, a marker/inducer of pluripotency and cellular clonogenicity [11, 19, 20]. Therefore, although we have not evaluated the clonogenic and differentiation capacity of rapamycin-treated BM-MSCs, the diminished secretion of IL6 and elevated expression of *NANOG* may confer stemness properties to rapamycin-treated BM-MSCs, such as the observed cell PD at a relatively more constant rate. However, the expression patterns of these two markers were not correlated with the rapamycin-mediated increase in the replicative capacity of each sample, suggesting that other molecular triggers are involved in this process.

Finally, our data show a significant correlation between rapamycin-mediated lifespan extension of BM-MSCs and inhibition of p16^INK4A^ protein expression, as more significant reductions in p16^INK4A^ accumulation during successive cellular passages were found in rapamycin-treated cells which showed the longer lifespans. Although previous studies using cultured fibroblast have shown that chromatin-remodeling patterns of *p16*^*INK4A*^ promoter region induced by caloric restriction (which is known to inhibits mTOR signaling) [21] or enhanced p16^INK4A^ mRNA decay mediated by RNA binding proteins mainly expressed in non-senescent cells [22] may account to inhibit p16^INK4A^ accumulation, our results suggest that the mechanisms underlying the rapamycin-mediated lifespan prolongation in long-term cultured BM-MSCs may involve either suppression of p16^INK4A^ mRNA translation or enhanced degradation of p16^INK4A^ protein, as p16^INK4A^ transcript levels were not decreased in rapamycin-treated BM-MSCs. Indeed, suppression of mTOR is well known to attenuate the translation of specific mRNAs [18] and to augment autophagy [23, 24], two effector programs of cellular senescence [20, 25].

It is noteworthy that despite the presence of rapamycin at a dose that attenuates mTOR signaling along the entire experiment, all rapamycin-treated samples progressed towards growth arrest and, with exception of BM12 that did not show any response to rapamycin, showed only a slight increased level of p16^INK4A^ expression at the final passage compared to the same cells at an intermediate passage (“deviation passage with rapamycin”), suggesting that mTOR- and p16^INK4A^-independent mechanisms may contribute to replicative senescence of late passage rapamycin-treated cells. p16^INK4A^, a cyclin-dependent kinase inhibitor, negatively regulates cell cycle progression by inhibiting transition from the G1 to S phase [26], and an aging-associated increase in its expression have been shown to contribute to the decline in replicative potential of various progenitor cell types [27–29]. Nevertheless, the mechanisms by which p16^INK4A^ are involved in cellular senescence, either as a cause of senescence or a consequence of it, remain to be explicitly elucidated [30].

Taken together, the results reported herein provide evidence supporting the potential use of rapamycin to delay senescence of BM-MSCs *in vitro* and the critical role of p16^INK4A^ regulation in this process. However, caution should be taken when using rapamycin for this purpose as interindividual variability in response to rapamycin exists for unknown reasons, but that may involve modulators of p16^INK4A^ protein levels. It is possible that other mTOR inhibitors or drugs that regulate p16^INK4A^ turnover could exert similar effects on MSC expansion which opens new avenues of pharmacologic modulation of longevity. Thus, further investigation to unravel the molecular mechanisms of lifespan promoting effects of rapamycin would be of importance for increase long-term expansion of MSCs with maintenance of important cellular properties.

## Acknowledgments

The authors are grateful to all of the subjects who participated in this work. CAP is a visiting scientist supported by an institutional grant from Sociedade Beneficiente Israelita Brasileira Albert Einstein.

## Supporting information

**Fig S1.**
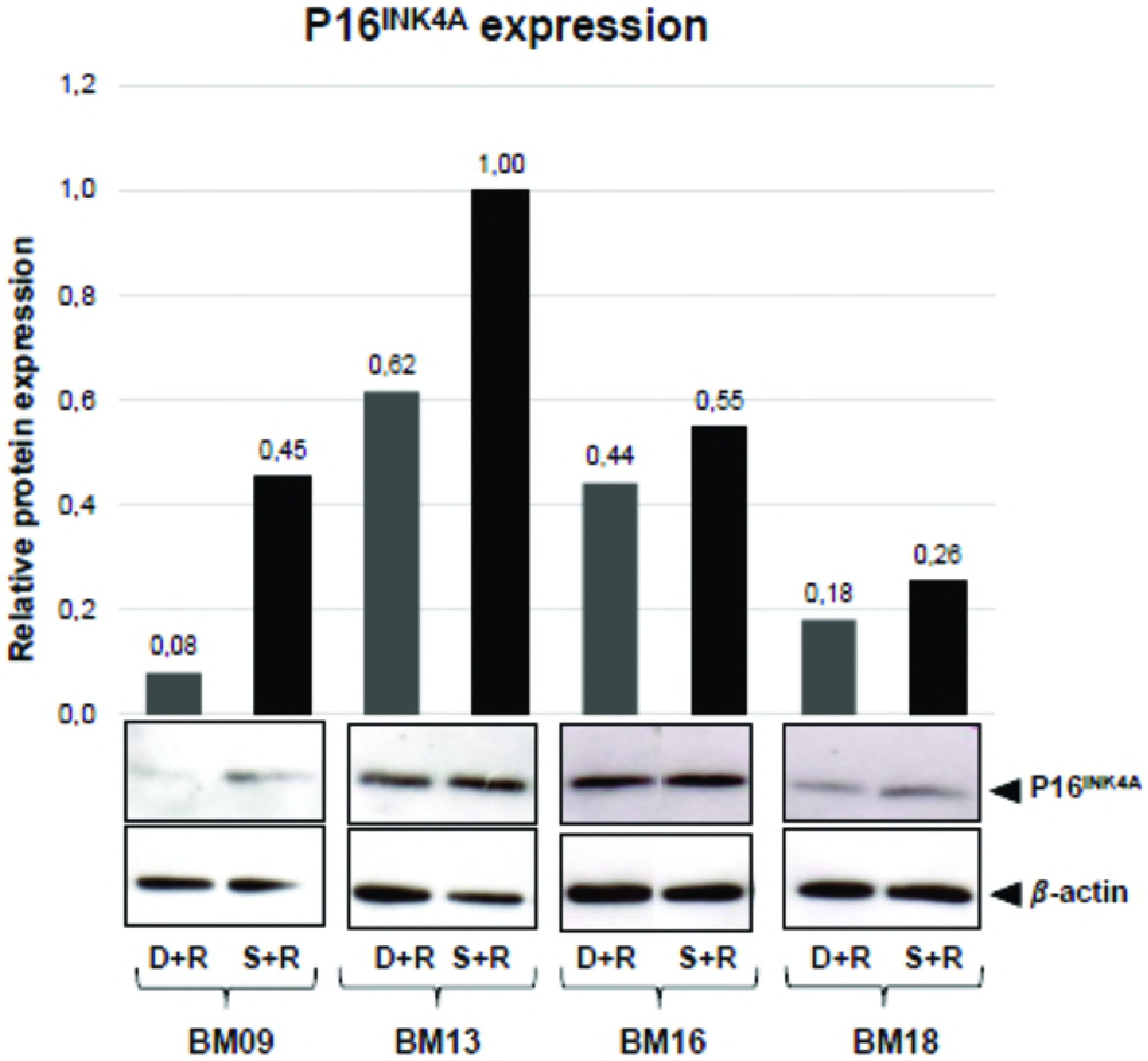
Expression levels of senescence- and pluripotency-related markers at an early passage as well as the replicative capacity of untreated BM-MSC samples were not correlated with the rapamycin-mediated replicative lifespan extension of BM-MSCs. The normalized expression values of senescence-associated proteins (p16^INK4^A, p21^WAF1^, pRPS6, SOD2, IL6 and IL8) and pluripotency-related genes (*NANOG, SOX2 and OCT4*) in each BM-MSC sample at early passages (5^th^ and 6^th^ passages) (previously determined in Piccinato et al., 2015) [14] **(A)**, as well the final PD number of each BM-MSC sample expanded in normal medium (without rapamycin) **(B)**, were plotted against the additional PD number obtained for the corresponding rapamycin-treated cells and statistically analyzed by Spearman correlation, as shown in the graphs. RAPA= rapamycin. PD= population doubling.

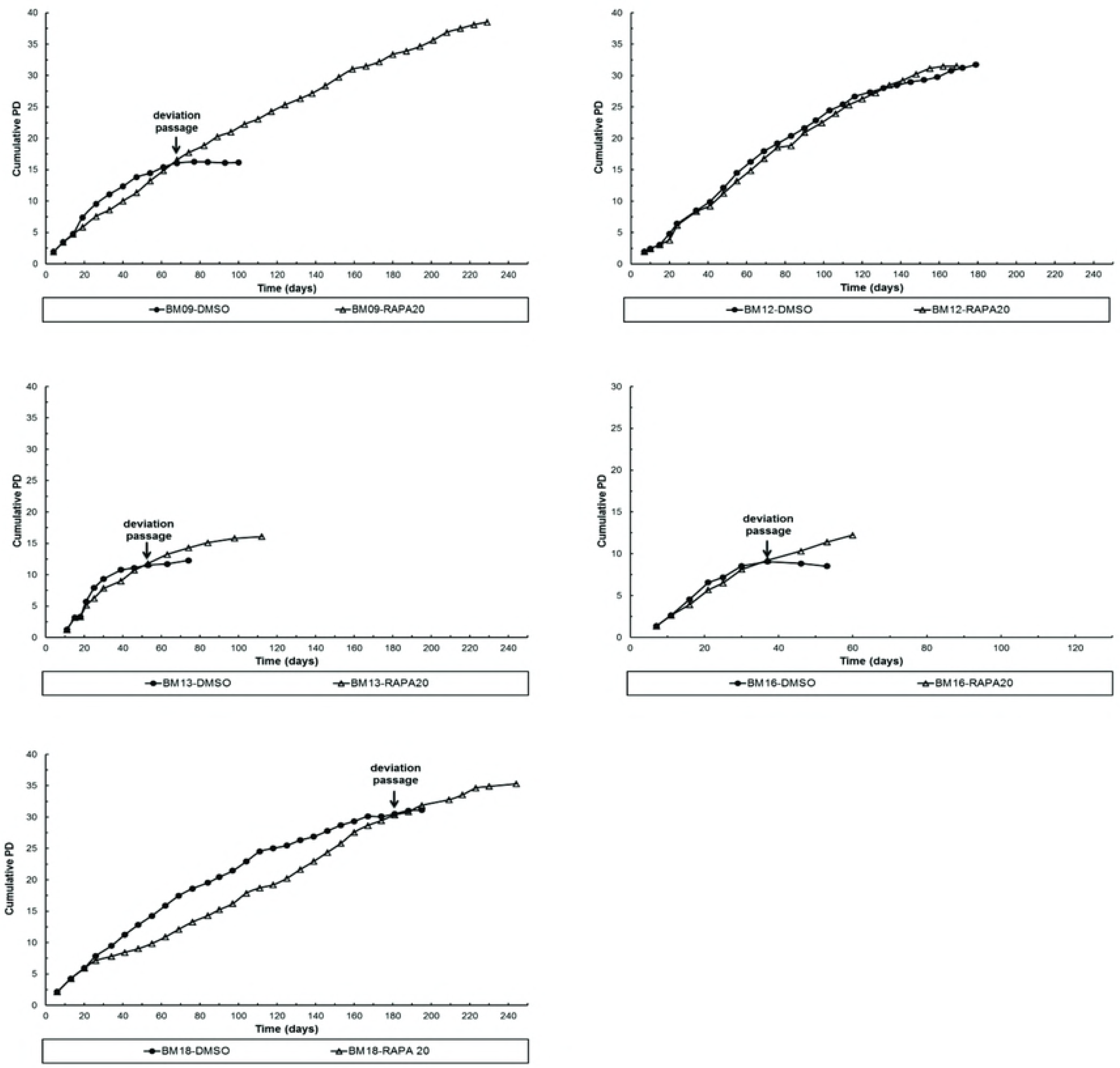

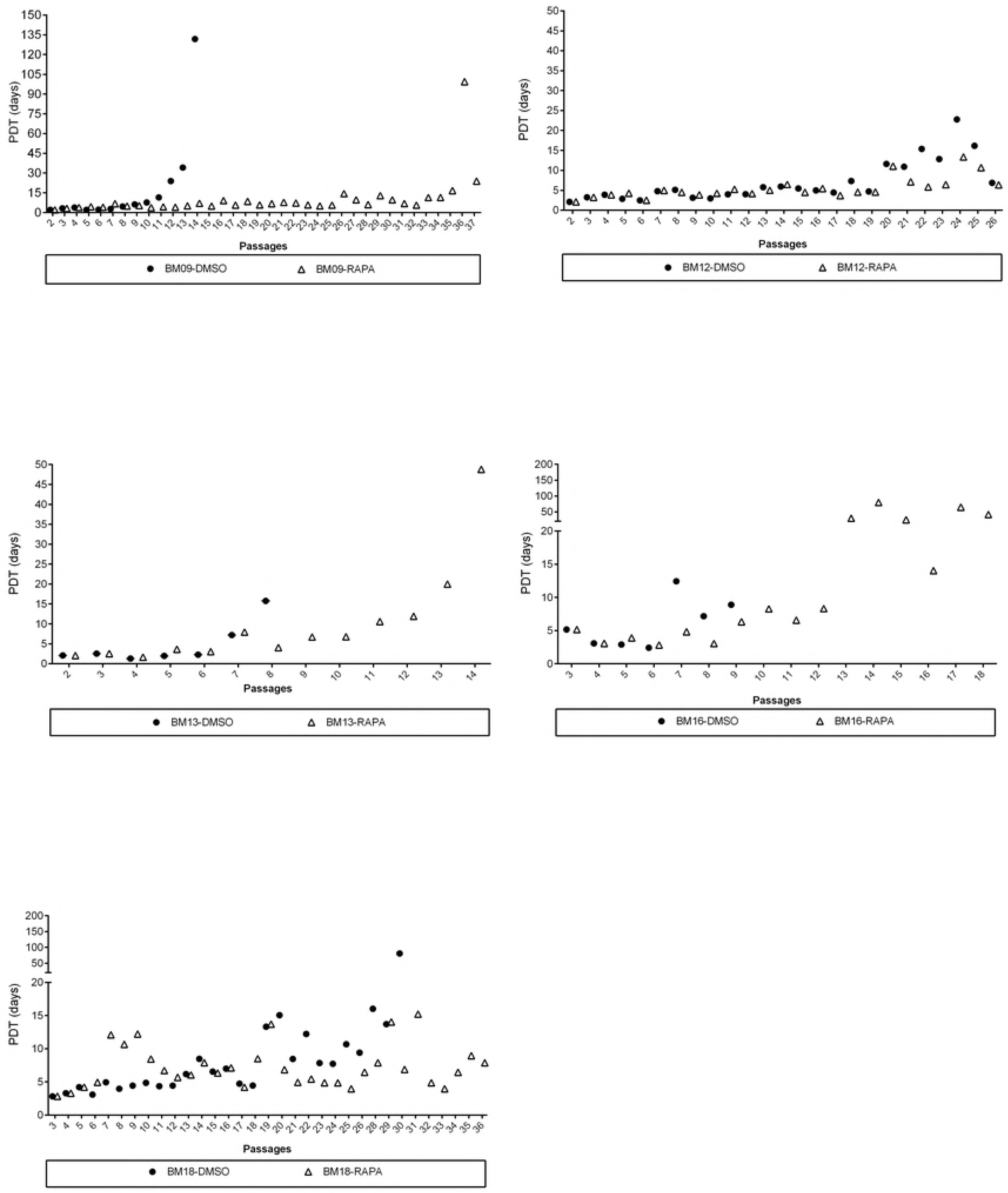

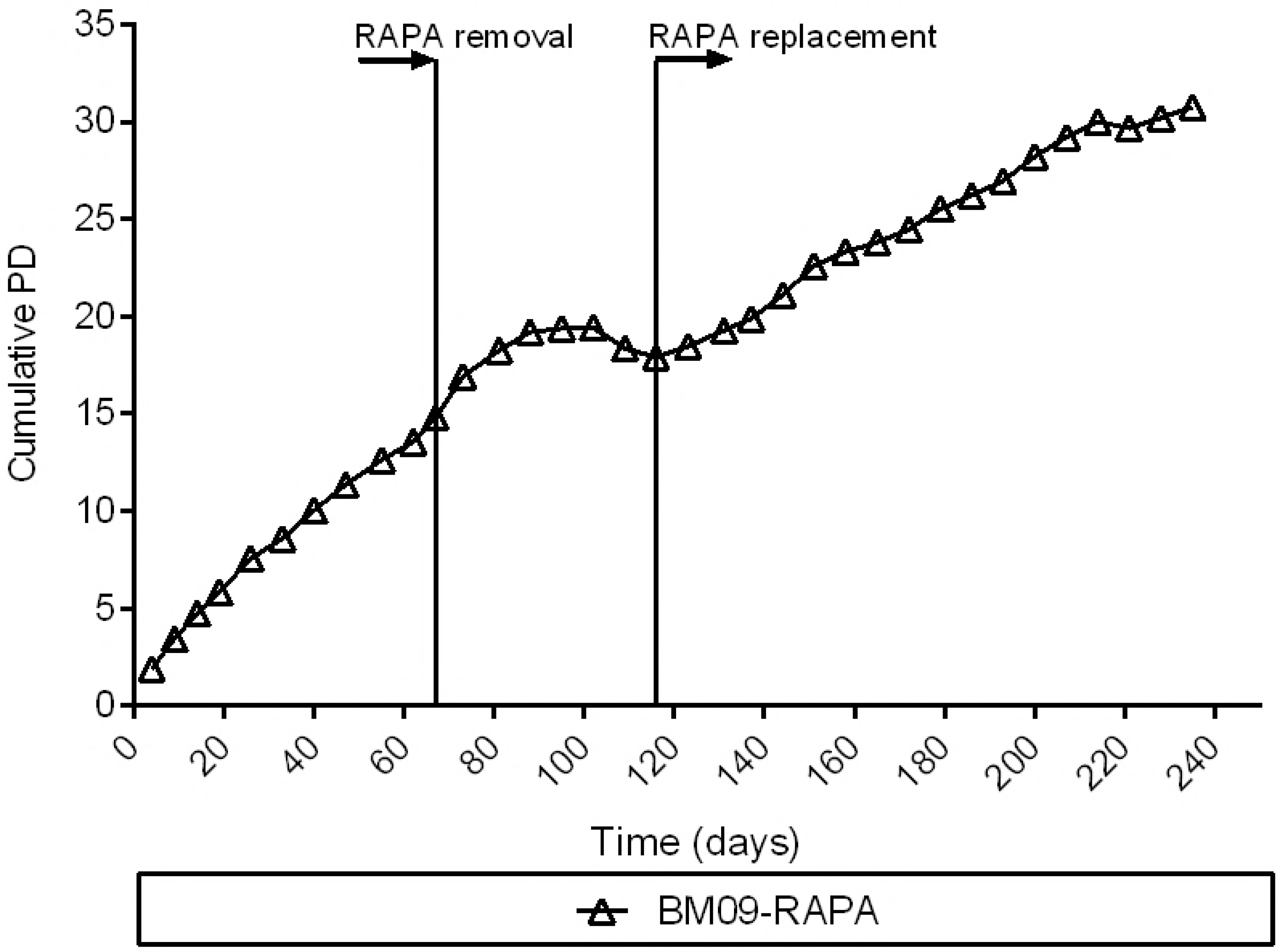

## References

1. Johnson SC, Rabinovitch PS, Kaeberlein M. mTOR is a key modulator of ageing and age-related disease. Nature. 2013;493(7432):338–45. doi: 10.1038/nature11861. PubMed PMID: 23325216; PubMed Central PMCID: PMC3687363.

2. Xu S, Cai Y, Wei Y. mTOR Signaling from Cellular Senescence to Organismal Aging. Aging and disease. 2014;5(4):263–73. doi: 10.14336/AD.2014.0500263. PubMed PMID: 25110610; PubMed Central PMCID: PMC4113516.

3. Saxton RA, Sabatini DM. mTOR Signaling in Growth, Metabolism, and Disease. Cell. 2017;169(2):361–71. doi: 10.1016/j.cell.2017.03.035. PubMed PMID: 28388417.

4. Harrison DE, Strong R, Sharp ZD, Nelson JF, Astle CM, Flurkey K, et al. Rapamycin fed late in life extends lifespan in genetically heterogeneous mice. Nature. 2009;460(7253):392–5. Epub 2009/07/08. doi: 10.1038/nature08221. PubMed PMID: 19587680; PubMed Central PMCID: PMCPMC2786175.

5. Chen C, Liu Y, Zheng P. mTOR regulation and therapeutic rejuvenation of aging hematopoietic stem cells. Sci Signal. 2009;2(98):ra75. Epub 2009/11/24. doi: 10.1126/scisignal.2000559. PubMed PMID: 19934433; PubMed Central PMCID: PMCPMC4020596.

6. Iglesias-Bartolome R, Patel V, Cotrim A, Leelahavanichkul K, Molinolo AA, Mitchell JB, et al. mTOR inhibition prevents epithelial stem cell senescence and protects from radiation-induced mucositis. Cell Stem Cell. 2012;11(3):401–14. doi: 10.1016/j.stem.2012.06.007. PubMed PMID: 22958932; PubMed Central PMCID: PMCPMC3477550.

7. Jang YY, Sharkis SJ. A low level of reactive oxygen species selects for primitive hematopoietic stem cells that may reside in the low-oxygenic niche. Blood. 2007;110(8):3056–63. Epub 2007/06/26. doi: 10.1182/blood-2007-05-087759. PubMed PMID: 17595331; PubMed Central PMCID: PMCPMC2018677.

8. Gan B, DePinho RA. mTORC1 signaling governs hematopoietic stem cell quiescence. Cell Cycle. 2009;8(7):1003–6. Epub 2009/04/02. doi: 10.4161/cc.8.7.8045. PubMed PMID: 19270523; PubMed Central PMCID: PMCPMC2743144.

9. Castilho RM, Squarize CH, Chodosh LA, Williams BO, Gutkind JS. mTOR mediates Wnt-induced epidermal stem cell exhaustion and aging. Cell Stem Cell. 2009;5(3):279–89. doi: 10.1016/j.stem.2009.06.017. PubMed PMID: 19733540; PubMed Central PMCID: PMCPMC2939833.

10. Hobbs RM, Seandel M, Falciatori I, Rafii S, Pandolfi PP. Plzf regulates germline progenitor self-renewal by opposing mTORC1. Cell. 2010;142(3):468–79. doi: 10.1016/j.cell.2010.06.041. PubMed PMID: 20691905; PubMed Central PMCID: PMCPMC3210556.

11. Gharibi B, Farzadi S, Ghuman M, Hughes FJ. Inhibition of Akt/mTOR attenuates age-related changes in mesenchymal stem cells. Stem Cells. 2014;32(8):2256–66. doi: 10.1002/stem.1709. PubMed PMID: 24659476.

12. Wagner W, Horn P, Castoldi M, Diehlmann A, Bork S, Saffrich R, et al. Replicative senescence of mesenchymal stem cells: a continuous and organized process. PLoS One. 2008;3(5):e2213. doi: 10.1371/journal.pone.0002213. PubMed PMID: 18493317; PubMed Central PMCID: PMC2374903.

13. Wagner W, Ho AD, Zenke M. Different facets of aging in human mesenchymal stem cells. Tissue engineering Part B, Reviews. 2010;16(4):445–53. doi: 10.1089/ten.TEB.2009.0825. PubMed PMID: 20196648.

14. Piccinato CA, Sertie AL, Torres N, Ferretti M, Antonioli E. High OCT4 and Low p16(INK4A) Expressions Determine In Vitro Lifespan of Mesenchymal Stem Cells. Stem Cells Int. 2015;2015:369828. Epub 2015/05/21. doi: 10.1155/2015/369828. PubMed PMID: 26089914; PubMed Central PMCID: PMCPMC4454755.

15. Blagosklonny MV. From rapalogs to anti-aging formula. Oncotarget. 2017;8(22):35492–507. doi: 10.18632/oncotarget.18033. PubMed PMID: 28548953; PubMed Central PMCID: PMC5482593.

16. Leontieva OV, Blagosklonny MV. Gerosuppression by pan-mTOR inhibitors. Aging (Albany NY). 2016;8(12):3535–51. doi: 10.18632/aging.101155. PubMed PMID: 28077803; PubMed Central PMCID: PMC5270685.

17. Wullschleger S, Loewith R, Hall MN. TOR signaling in growth and metabolism. Cell. 2006;124(3):471–84. doi: 10.1016/j.cell.2006.01.016. PubMed PMID: 16469695.

18. Laberge RM, Sun Y, Orjalo AV, Patil CK, Freund A, Zhou L, et al. MTOR regulates the pro-tumorigenic senescence-associated secretory phenotype by promoting IL1A translation. Nat Cell Biol. 2015;17(8):1049–61. Epub 2015/07/06. doi: 10.1038/ncb3195. PubMed PMID: 26147250; PubMed Central PMCID: PMCPMC4691706.

19. Pospelova TV, Bykova TV, Zubova SG, Katolikova NV, Yartzeva NM, Pospelov VA. Rapamycin induces pluripotent genes associated with avoidance of replicative senescence. Cell Cycle. 2013;12(24):3841–51. Epub 2013/12/02. doi: 10.4161/cc.27396. PubMed PMID: 24296616; PubMed Central PMCID: PMCPMC3905076.

20. Rubinsztein DC, Marino G, Kroemer G. Autophagy and aging. Cell. 2011;146(5):682–95. doi: 10.1016/j.cell.2011.07.030. PubMed PMID: 21884931.

21. Li Y, Tollefsbol TO. p16(INK4a) suppression by glucose restriction contributes to human cellular lifespan extension through SIRT1-mediated epigenetic and genetic mechanisms. PLoS One. 2011;6(2):e17421. Epub 2011/02/24. doi: 10.1371/journal.pone.0017421. PubMed PMID: 21390332; PubMed Central PMCID: PMCPMC3044759.

22. Wang W, Martindale JL, Yang X, Chrest FJ, Gorospe M. Increased stability of the p16 mRNA with replicative senescence. EMBO Rep. 2005;6(2):158–64. doi: 10.1038/sj.embor.7400346. PubMed PMID: 15678155; PubMed Central PMCID: PMCPMC1299256.

23. Jung CH, Jun CB, Ro SH, Kim YM, Otto NM, Cao J, et al. ULK-Atg13-FIP200 complexes mediate mTOR signaling to the autophagy machinery. Mol Biol Cell. 2009;20(7):1992–2003. Epub 2009/02/18. doi: 10.1091/mbc.E08-12-1249. PubMed PMID: 19225151; PubMed Central PMCID: PMCPMC2663920.

24. Lerner C, Bitto A, Pulliam D, Nacarelli T, Konigsberg M, Van Remmen H, et al. Reduced mammalian target of rapamycin activity facilitates mitochondrial retrograde signaling and increases life span in normal human fibroblasts. Aging cell. 2013;12(6):966–77. doi: 10.1111/acel.12122. PubMed PMID: 23795962; PubMed Central PMCID: PMC5559196.

25. Hands SL, Proud CG, Wyttenbach A. mTOR’s role in ageing: protein synthesis or autophagy? Aging (Albany NY). 2009;1(7):586–97. doi: 10.18632/aging.100070. PubMed PMID: 20157541; PubMed Central PMCID: PMC2806042.

26. Li J, Poi MJ, Tsai MD. Regulatory mechanisms of tumor suppressor P16(INK4A) and their relevance to cancer. Biochemistry. 2011;50(25):5566–82. doi: 10.1021/bi200642e. PubMed PMID: 21619050; PubMed Central PMCID: PMC3127263.

27. Krishnamurthy J, Ramsey MR, Ligon KL, Torrice C, Koh A, Bonner-Weir S, et al. p16INK4a induces an age-dependent decline in islet regenerative potential. Nature. 2006;443(7110):453–7. doi: 10.1038/nature05092. PubMed PMID: 16957737.

28. Molofsky AV, Slutsky SG, Joseph NM, He S, Pardal R, Krishnamurthy J, et al. Increasing p16INK4a expression decreases forebrain progenitors and neurogenesis during ageing. Nature. 2006;443(7110):448–52. doi: 10.1038/nature05091. PubMed PMID: 16957738; PubMed Central PMCID: PMC2586960.

29. Janzen V, Forkert R, Fleming HE, Saito Y, Waring MT, Dombkowski DM, et al. Stem-cell ageing modified by the cyclin-dependent kinase inhibitor p16INK4a. Nature. 2006;443(7110):421–6. doi: 10.1038/nature05159. PubMed PMID: 16957735.

30. Coppe JP, Rodier F, Patil CK, Freund A, Desprez PY, Campisi J. Tumor suppressor and aging biomarker p16(INK4a) induces cellular senescence without the associated inflammatory secretory phenotype. The Journal of biological chemistry. 2011;286(42):36396–403. doi: 10.1074/jbc.M111.257071. PubMed PMID: 21880712; PubMed Central PMCID: PMC3196093.

